# Continuous two-phase *in vitro* co-culture model of the enthesis

**DOI:** 10.1101/2022.02.07.479445

**Authors:** Hyeree Park, Megan E. Cooke, Jean-Gabriel Lacombe, Michael H. Weber, Paul A. Martineau, Showan N. Nazhat, Derek H. Rosenzweig

**Affiliations:** Department of Mining and Materials Engineering, McGill University, Montreal, Canada; Department of Experimental Surgery, McGill University, Montreal, Canada; Injury, Repair and Recovery Program, Research Institute of McGill University Health Centre, Montreal, Canada; Division of Orthopaedic Surgery, McGill University, Montreal, Canada

**Keywords:** dense collagen, hydrogel, tendon, ligament, alignment, anisotropy, bone, *in vitro* model

## Abstract

The enthesis is the interfacial tissue between ligament or tendon, and bone, which connects tissues of distinctly different mechanical properties. Although ligament and enthesis injury is commonplace, the development and healing mechanisms of these tissues are yet unclear. Current models for investigating these mechanisms are primarily *in vivo* animal models as *in vitro* models have been limited. In this study, an *in vitro* enthesis model was developed using a modified gel aspiration-ejection (GAE) method. Continuous two-phase aligned dense collagen (ADC) hydrogels with an overlapping interface were fabricated within 2 hours. The mechanical properties of acellular two-phase ADC confirmed the continuous nature of this model, as the mechanical properties showed no significant difference compared to single-phase ADC and maintained comparable structural and mechanical characteristics of immature ligaments and unmineralized bone. Human anterior cruciate ligament (ACL) fibroblasts and human spine vertebral osteoblasts were isolated from donor tissues and were seeded to form the enthesis model. These were cultured for 14 days, at which the viability and proliferation was observed to be 85 ± 7.5% and 230 ± 52%, respectively. Histological and immunofluorescence analyses at day 14 revealed extensive cell-driven matrix remodelling, and the seeded fibroblasts and osteoblasts maintained their phenotype within their compartments of the layered co-culture model. These results demonstrate the rapid fabrication of a two-phase co-culture system that can be utilized as an *in vitro* model to further understand the degenerative and regenerative mechanisms within the enthesis.

## 1. Introduction

The enthesis is the interface between ligament or tendon and bone, comprised of a matrix of dense tissue that functions as an arrested and “miniature” growth plate with a distinct morphological gradient of fibrocartilage between ligament and bone [1]. It facilitates load bearing and force transmitting functions whilst maintaining the connection between ligament, a soft tissue, and bone, a hard tissue. Approximately 33 million musculoskeletal injuries are reported annually in the United States alone, wherein approximately 50% of these injuries affect ligaments and tendons [2]. Injuries to these connective tissues and the enthesis are associated with pain and reduced range of motion that can be debilitating. Spinal instability secondary to ligamentous disruption can also result in progressive deformity and devastating neurological injuries. Due to the minimal vasculature present, healing is limited and the scar formed can be prone to further injury [3,4]. Current strategies for treating these diseases and injuries largely revolve around pain management, immobilization and often surgical intervention [5,6], with an estimated 300,000 tendon and ligament surgeries are performed annually in the United States alone [7]. Surgical reconstruction of the anterior cruciate ligament (ACL) surgeries can result in reduced biomechanical function and surgeries using autograft and allografts have demonstrated a failure rate of 11.9% over a 10-year period [8]. Despite the high prevalence of injury, replacement surgeries, and their subsequent failures, the mechanisms that affect the healing of ligament and enthesis after injury are poorly understood.

To investigate mechanisms that drive certain biological functions, both *in vivo* and *in vitro* models can be employed. For musculoskeletal tissues, *in vivo* models are preferred and remain the gold standard in preclinical studies due to a lack of reliable and biomimetic *in vitro* models [9]. *In vivo* models have been crucial in providing insight into enthesis development pathways [10,11], but a limited number of *in vitro* models have been developed [12–14]. Driven by the high occurrence of reconstruction surgeries for ligament and tendon tissues, there has been increased interest in providing bi- and tri-phasic scaffolds as alternative grafts, including differentiating mesenchymal stem cells into two lineages within the same scaffold [13,15–18]. A 2D zonal co-culture model between bone fibroblasts and osteoblasts with an overlapping interface has been proposed [14], but it is widely accepted that 3D scaffolds provide a more physiological microenvironment for cells [19]. In a recent study to develop a 3D model for fibroblast and osteoblast co-culture, various hydrogels including agarose, gellan, fibrin and collagen were investigated [12]. These hydrogels were seeded with chick tendon fibroblasts and murine pre-osteoblasts in various orientations with the aid of silicone moulds. The agarose and gellan scaffolds limited the adherence of the cells, whereas the highly hydrated collagen scaffolds were mechanically unstable when the mould was removed. On the other hand, fibrin scaffolds promoted cellular alignment of the seeded fibroblasts, which was driven by gravity to migrate towards adherent plates [12]. Importantly, the enthesis and its attached tissues are mechanoresponsive and require mechanical stimulation for development and healing. Therefore, there is a need to provide biomimetic *in vitro* models that more accurately depict the enthesis and where biochemical and mechanical stimulation can be applied to observe their effects.

Collagen type I is the most abundant protein in the human body, and in ligament, bone and their enthesis [20]. As such, collagen type I is a widely studied biomimetic scaffold material [21].

Neutralized collagen hydrogel concentrations are often between 2-6 mg/mL, and form highly hydrated collagen gels of randomly oriented fibrils [21], which are mechanically unstable [12]. By manipulating these highly hydrated collagen hydrogels, the collagen fibrils can be densified and aligned to form scaffolds that mimic the properties of immature musculoskeletal tissues [22–26]. By applying pressure differentials through a needle connected to an angioplasty inflation device to the highly hydrated collagen gels, simultaneous densification and alignment of the collagen fibers has been achieved. This gel aspiration-ejection (GAE) method aspirates the gel into the needle and removes the excess casting fluid, during which the shear forces that are applied to the gel aligns the collagen fibrils. Aligned dense collagen (ADC) scaffolds have previously been investigated *in vitro* seeded with single cell types, including fibroblasts, mesenchymal stem cells, vascular smooth muscle cells, and Schwann cells [21,24–26], and as a co-culture model using Schwann cells and neuronal cells [25]. The latter, however, did not form zones for the co-culture but seeded secondary cells on the scaffold surface. In this study, the existing GAE method was modified to form a continuous two-phase ADC scaffold with two distinct but interfacing zones. This model was initially validated using a collagen-binding dye and fluorescent expressing cells as proof-of-concept. The enthesis model was then fabricated by seeding ADC gels with primary human ACL fibroblasts and osteoblasts, which were cultured for 14 days. Cell viability, proliferation as well as the cell-driven matrix remodelling was observed. This study proposes and demonstrates the fabrication of a continuous co-culture system that can be used as an *in vitro* tool to investigate the enthesis.

## 2. Experimental Methods

### 2.1 Cell isolation and culture

Fluorescent-protein expressing cell-lines, GFP-MDA-MB-231 adenocarcinoma and mCherry-IRM90 lung fibroblasts were cultured in T75 flasks in basal media, high-glucose Dulbecco’s modified Eagle medium (DMEM, Gibco, Canada) supplemented with 10% fetal bovine serum (Gibco, Canada) and 1% penicillin/streptomycin (P/S, Gibco, Canada). Cells were passaged at 80% confluency and were used up to passage 6.

Human ACL fibroblasts were isolated from tissues obtained from patients undergoing ACL reconstruction surgeries. Human ACL tissues were collected with consent and approved by the ethical review board at the McGill University (IRB A04-M53-08B and A10-M113-13B). The tissues were washed in 2X phosphate buffer solution (PBS) supplemented with 2% Amphotericin B (Gibco, Canada) and 2% P/S, and digested for 16 h at 37 °C in 1.5 mg/mL collagenase type II solution (Gibco, Canada) in basal media. The digested tissues were strained in 100 and subsequently 70 µm cell strainers and plated in adherent T75 flasks. Cells were passaged at 80% confluency and were used from passage 1-4. Human osteoblasts were isolated from cortical bone from spinal processes [27]. The collection of human lumbar spinal tissue was approved by the ethical review board at Research Institute of the Montreal University Health Center (RI-MUHC, REB Extracellular Matrix Protocol # 2020-5647) and obtained with informed consent in collaboration with Transplant Quebec. After the physical removal of soft tissues, bone chips of approximately 3-5 mm^3^ were cut from the spinal processes and washed with 2X PBS supplemented with antibiotics. To further remove any remaining soft tissues, the chips were incubated with collagenase type II solution as above. The bone chips were re-washed and plated in T75 flasks. After cells reached approximately 30 – 40% confluency from the outgrowth of osteoblasts, the bone chips were removed from the T75 flasks. Cells were passaged at 80% confluency and were used from passage 1-4. Demographics of each ACL fibroblast-spinal osteoblast co-culture is detailed in Table 1.

**Table 1.**
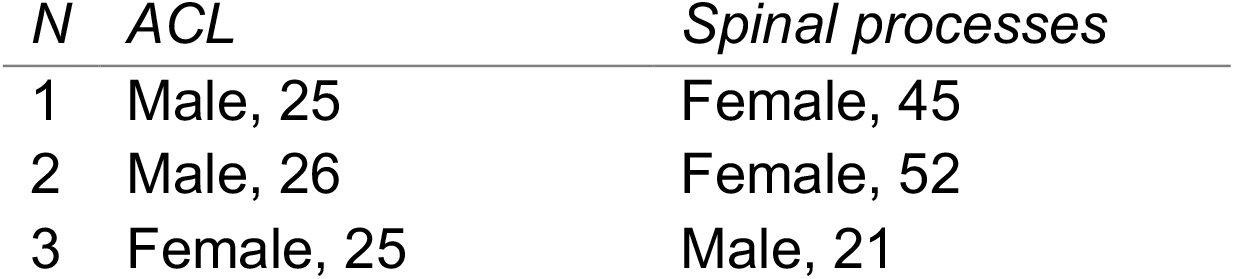
Tissue donor demographics indicating age and sex

### 2.2 Fabrication of acellular and cellular two-phase aligned dense collagen scaffolds

Continuous two-phase ADC hydrogels were fabricated using the GAE method as previously described [23] with minor modifications (Figure 1). Acellular two-phased precursor highly hydrated collagen gels were first prepared using two independent collagen solutions, where 2.05 mg/mL rat-tail derived collagen type I (First Link, Ltd., UK) and 10x DMEM (First Link Ltd., UK) were mixed at a 4:1 volume ratio and neutralized with 5 M NaOH (Sigma, Canada). Fast Green (Sigma, Canada) was added at 0.02% to one of the neutralized collagen solutions (Figure 1A I). Aliquots of 500 µL of each of the two solutions were layered in 48-well plates (Figure 1A II inset). The layered collagen solutions underwent gelling at 37 °C for 30 minutes to form highly hydrated collagen gels (Figure 1A II). Once the two-phase highly hydrated gels were formed, an angioplasty inflation device (B. Braun, Germany) fitted with a 12-gauge (G) needle was for GAE, fabricating meso-scale anisotropic ADC gels.

Cellular two-phase ADCs were prepared by two solutions with 8:1:1 volume ratio of 2.05 mg/mL collagen type I, 10x DMEM, and cell-laden basal media, respectively. Two types of two-phase cell-seeded highly hydrated collagen gels were prepared: as proof-of-concept, GFP-MDA-231 (bottom) and mCherry-IRM90 lung fibroblasts (top) (Figure 1B); and for enthesis modelling, human ACL fibroblasts (bottom) and human spinal osteoblasts (top) (Figure 1C). Collagen solution and 10x DMEM were initially mixed and neutralized, and cell-laden basal media was subsequently added at a cellular density of 200,000 cells/mL pre-densification. Aliquots of 1 mL of each solution was cast in a 4 mL Wheaton Omni-Vial® (DWK Life Sciences, USA). Human ACL fibroblast-osteoblast seeded ADCs were cultured for 14 days tethered, *i*.*e*. clamped *in situ*, in a custom polycaprolactone mould designed for a 6-well plate (Figure 1C IV) [26] in basal medium supplemented with 50 µg/ml ascorbic acid to promote extracellular matrix formation [28]. As controls, single-phase ligament and osteoblast scaffolds were fabricated and cultured as above.

**Figure 1.**
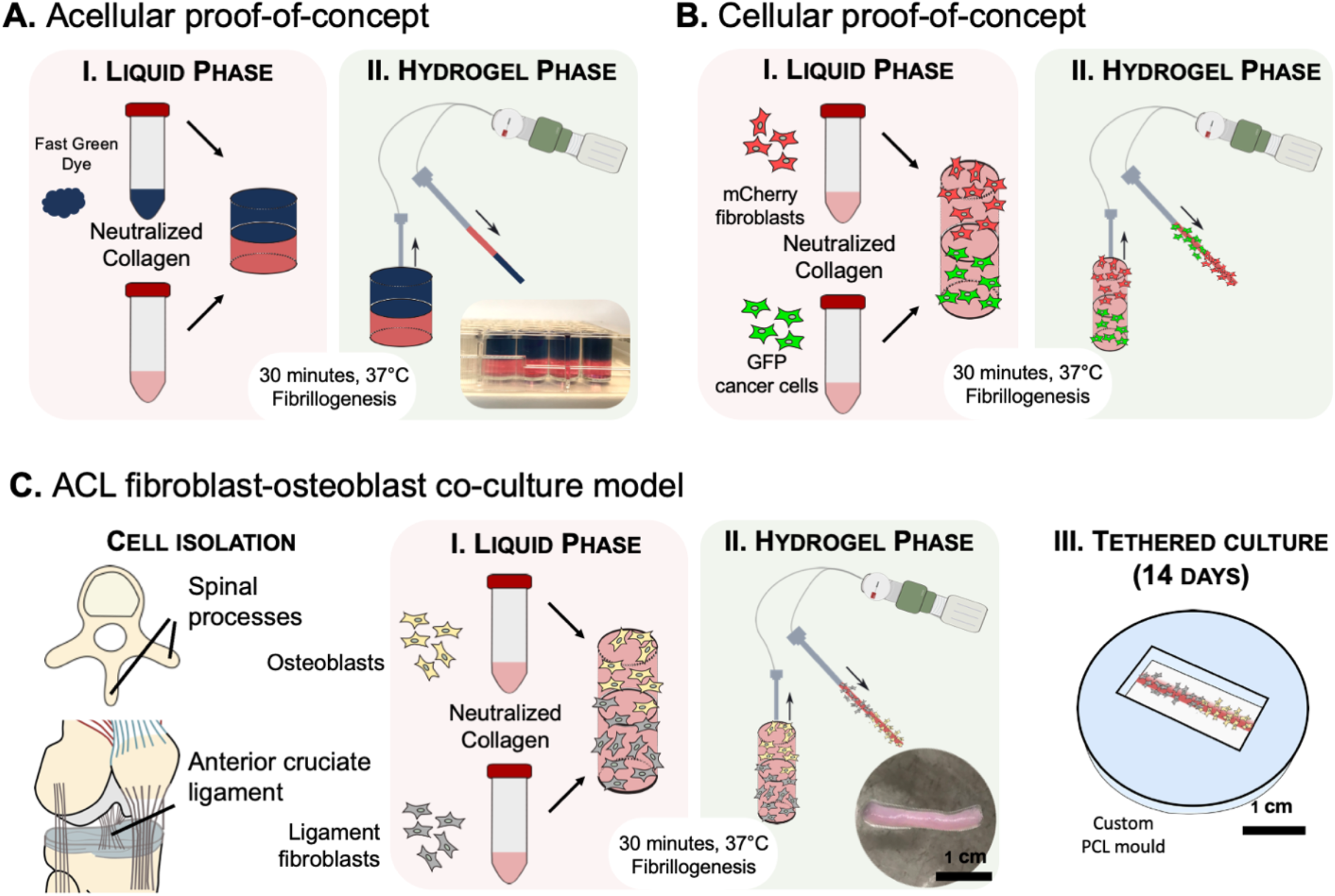
Gel aspiration-ejection (GAE) method to fabricate two-phased aligned dense collagen (ADC) scaffolds. Two independent collagen solutions were prepared and cast within a mould (I), which formed a highly hydrated precursor collagen hydrogel at 37 °C for 30 minutes. The precursor hydrogel was then aspirated and ejected using an angioplasty inflation device fitted with a 12G needle forming ADC scaffolds (II). (A) Acellular proof-of-concept two-phase scaffolds were fabricated using two neutralized collagen solutions, one of which was supplemented with 0.02% fast green dye (inset: image of precursor hydrogels), (B) Cellular two-phase scaffolds were fabricated by seeding GFP expressing MDA-MB-231 adenocarcinoma or mCherry expressing IRM90 lung fibroblasts into the two neutralized collagen solutions.(C) Enthesis co-culture model was fabricated using patient-isolated cells: ligament fibroblasts from ACLs and osteoblasts from spinal processes. The cells were seeded in individually neutralized collagen solutions (I), which underwent the GAE method to form a continuous two-phase hydrogel (inset) (II). The two-phase co-culture model was tethered within a mould for 14 days (III).

### 2.3 Aligned dense collagen hydrogel characterization

ADC gels were morphologically characterized by non-linear scanning microscopy [23]. Acellular ADC hydrogels were fixed in 4% formaldehyde and embedded in paraffin blocks and sectioned to 60 µm. The sections were imaged using a Leica SP8 MP (Leica, Germany) multiphoton microscope equipped with a 63x glycerin immersion objective with the laser excitation wavelength at 830 nm. Second harmonic generation emission was captured. Photographic images of the acellular two-phase ADCs were captured with a digital camera (Canon 360D). Cellular two-phase ADCs seeded with fluorescent expressing IRM90 (mCherry) and MDA-MB-231 (GFP) were observed at 2x using EVOS M5000 (ThermoFisher, Canada) through the GFP and Texas Red channels. Tensile tests of acellular ADCs were carried out (n = 5) using a 10 N load cell at a displacement rate of 0.1 mm/s (Univert Mechanical Tester, CellScale Biomaterials, Canada). The cross-sectional area of ADC was assumed to correspond to the internal diameter of the needle used, *i*.*e*. 2.16 mm for the 12G needle.

### 2.4 Cell viability and number

At day 14 of culture, LIVE/DEAD™ Viability/Cytotoxicity Kit (ThermoFisher, Canada) was used according to manufacturer’s instructions. Three images of three independent experiments (n = 3) of single-culture and co-culture ACL fibroblast-osteoblast scaffolds were captured using a 10x objective attached to EVOS M5000 (ThermoFisher, Canada). The percentage of live cells were quantified using the cell counter plug-in on ImageJ (NIH, USA). Total cell number was obtained by Hoechst 33258 DNA assay, as previously described [29,30]. Briefly, ADC scaffolds (n = 3) were gently agitated in 4 M guanidine hydrochloride buffer (GuHCl, Sigma-Aldrich, Canada) supplemented with a protease inhibitor (Roche Applied Science, USA) for 48 h at 4 °C. 1 µg/mL Hoechst 233258 (Invitrogen, USA) was prepared and the samples were analyzed in a 96-well microplate using the Tecan M200 plate reader, at 360 nm excitation and 460 nm emission, against serial dilutions of calf-thymus DNA (Invitrogen, USA) that provided a standard curve. The weight of single cell DNA was assumed to be 7 picograms [27].

### 2.5 Histological analysis

Two-phase fibroblast-osteoblast ADC scaffolds (n = 3) were fixed in 4% paraformaldehyde (Sigma, Canada) for 1 hour and submerged in gradient sucrose solutions from 10% to 30%. These samples were embedded in O.C.T. compound (TissueTek, Canada), snap frozen at -80 °C and cut to 7 µm sections using a CM1950 cryostat (Leica biosystems, Germany). The sections were stained with safranin O/fast green (Sigma-Aldrich, Canada) and imaged with a Zeiss Axioskop 40 microscope (Carl Zeiss, Germany). Immunofluorescence staining was also performed. Sections were permeabilized and blocked (5% bovine serum albumin, 0.1% Triton-X 100 in PBS) for 45 minutes and incubated overnight at 4 °C with one or two primary antibodies from different hosts. Primary antibodies used and their concentrations were as follows: collagen I (1:200, ab34710, Abcam, Canada), tenomodulin (1:200, ab203676, Abcam, Canada), scleraxis (1:200, ab58655, Abcam, Canada) and osteocalcin (OCN, 1:200, ab13418, Abcam, Canada). These were incubated for 1.5 h at room temperature with AlexaFluor 488 donkey anti-rabbit IgG (1:250, A21206, Invitrogen, Canada) and/or AlexaFluor 555 Goat anti-Mouse IgG (1:250, A21422, Invitrogen, Canada). Immunofluorescence images were captured using an EVOS M500 (ThermoFisher, Canada).

### 2.6 Statistical analysis

Experimental data was analysed using Graphpad Prism™ v9.0 (Graphpad Prism Software, USA.) Groups were analysed with unpaired Student t-tests and where applicable two-way ANOVA with multiple comparisons (Tukey’s).

## 3. Results

### 3.1 Continuous two-phase aligned dense collagen hydrogels

The successful fabrication of two-phase ADC scaffolds was demonstrated using both acellular and cellular methods. By adding fast green, a collagen-binding dye, to the top half of the pre-gelation solution (Figure 1B), a continuous two-phase ADC scaffold was fabricated where no material partition was observed, except for the gradient change in colour (Figure 2A). The acellular two-phase scaffold was characterized by non-linear scanning microscopy, which demonstrated collagen fibrillar alignment and scaffold anisotropy (Figure 2B). The tensile properties of the scaffolds were also investigated to confirm the continuous nature of the hydrogel (Figure 2C-D). The tensile strength was found to be 0.13 ± 0.05 MPa and apparent modulus was 0.54 ± 0.2 MPa, which were not significantly different from previous single-phase findings (p = 0.75 and 0.78, respectively) [26]. To confirm the two-phased nature of seeded cells within this modified GAE method, the distribution of fluorescent protein expressing cells were investigated in the two-phased ADC hydrogels. Fluorescence microscopy images confirmed that the mCherry expressing IRM90 and GFP expressing MDA-MB-231 cells were reorganized into two discrete areas with an overlapping interface between the two cell types (Figure 2E).

**Figure 2.**
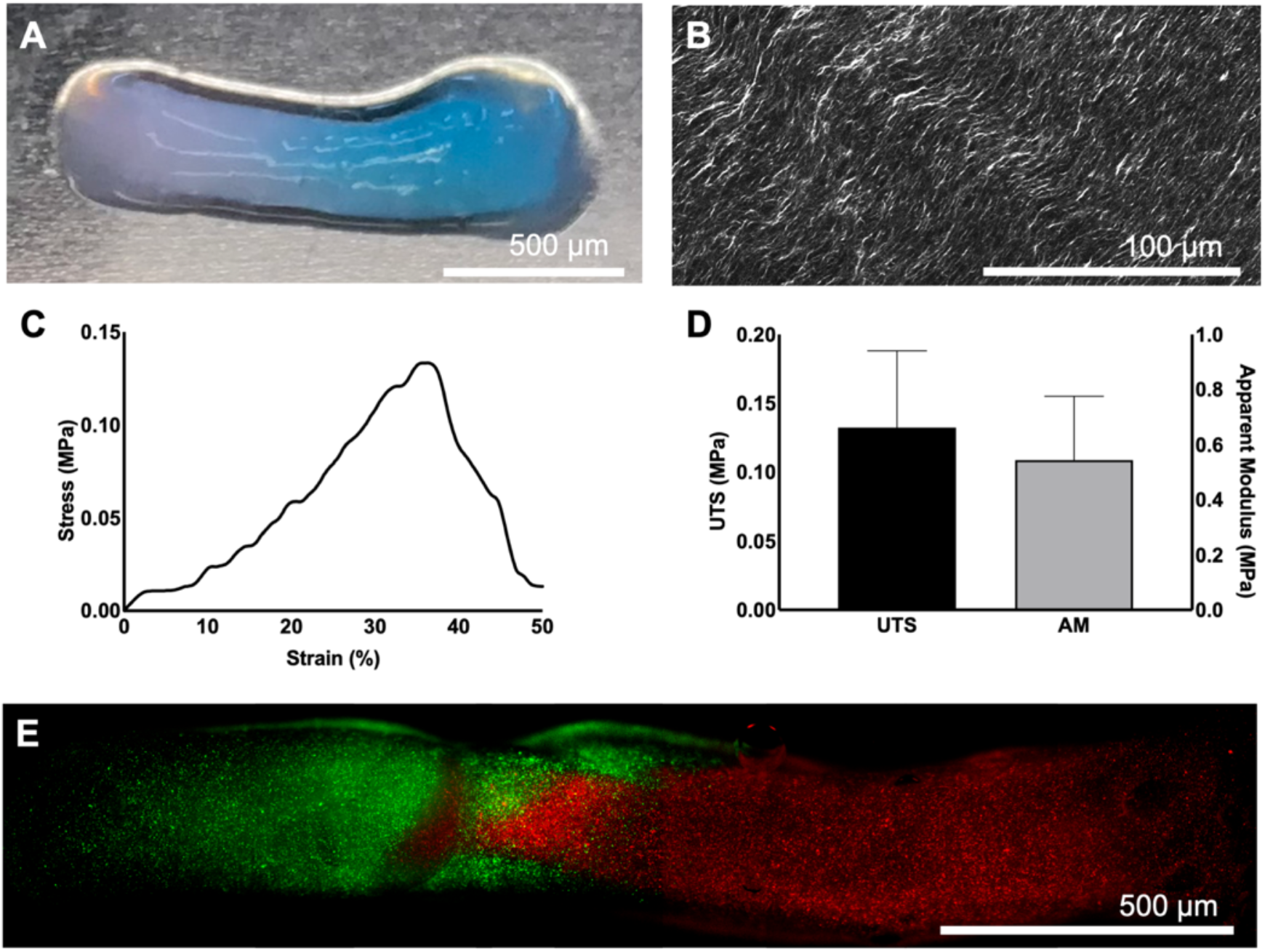
Two-phase ADC hydrogels. (A) Acellular two-phase ADC hydrogels with one phase dyed with Fast Green. (B) Non-linear scanning microscopy image of the acellular ADC hydrogel confirming their fibrillar alignment. (C) Representative tensile stress-strain curve for a two-phase ADC and (D) ultimate tensile stress (UTS) and apparent modulus (AM) of the scaffolds. Error bars ± SD. (E) Cell-seeded two-phase ADC gels as indicated by GFP expressing MDA-MB-231 (green) and mCherry expressing fibroblasts (red).

### 3.2 ACL fibroblast-osteoblast model

ACL fibroblast-osteoblast seeded ADC scaffolds were fabricated and cultured by tethering in custom moulds and cultured for 14 days (Figure 1C III) [26]. The percentage of live cells within these co-culture scaffolds was 85 ± 7.5%, which was not significantly different compared to their single-culture controls (p > 0.05) (Figure 3A and B). The proliferation of cells within the scaffolds was also investigated through a DNA assay, where high proliferative rates were observed (Figure 3C). The co-culture scaffolds proliferated by 230 ± 52%, whereas the single fibroblast and osteoblast cultured scaffolds by 189 ± 61% and 333 ± 58%, respectively. The osteoblast-seeded scaffolds showed significantly (p = 0.048) higher proliferation rates than the fibroblast-seeded scaffolds.

**Figure 3.**
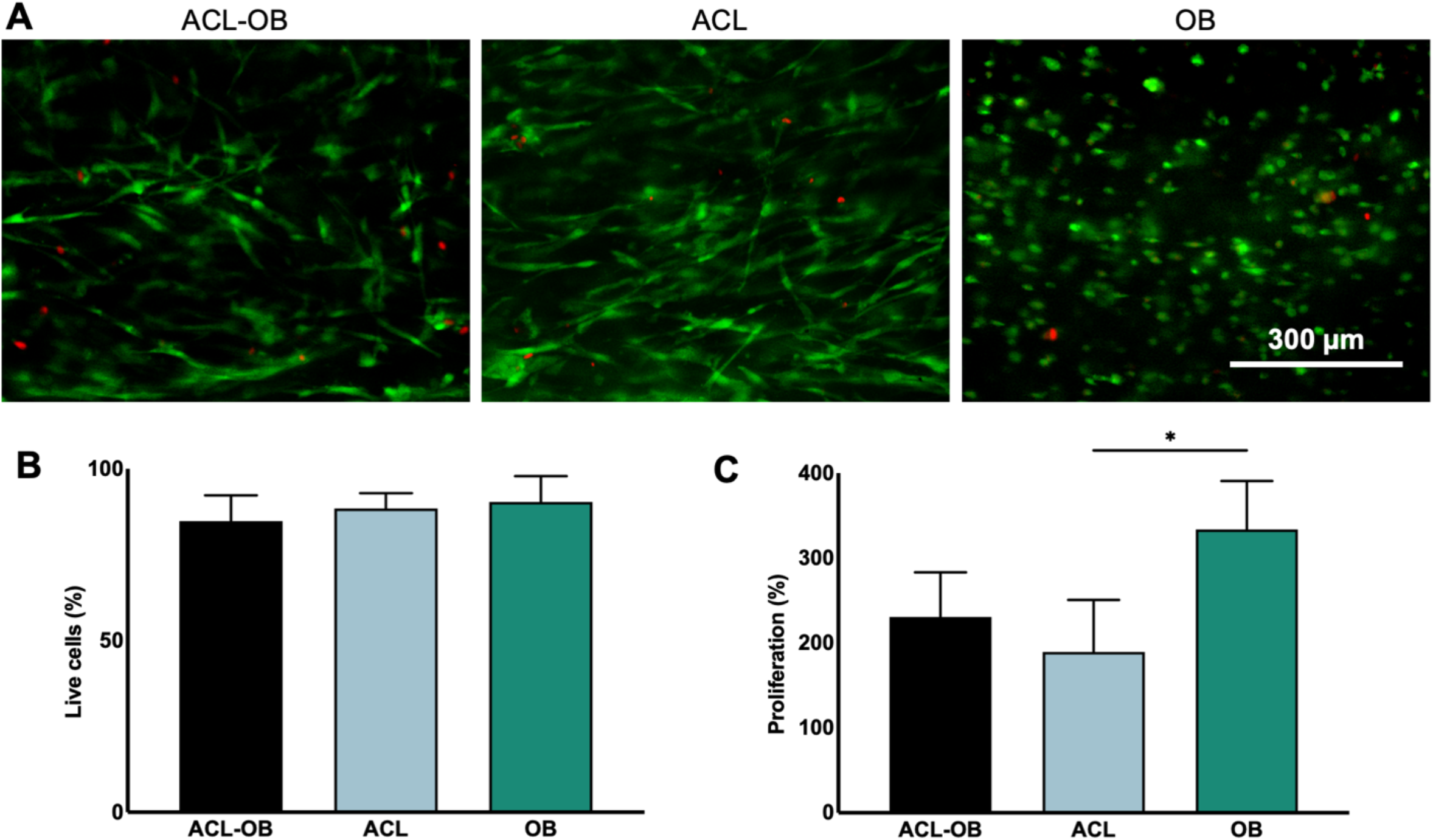
Viability and proliferation of ACL fibroblast-osteoblast co-culture model in ADC scaffolds at day 14. (A) Representative images of LIVE/DEAD of ACL fibroblast-osteoblast-seeded scaffolds and their single-culture controls. (B) Quantification of the LIVE/DEAD images. (C) Percentage proliferation of the seeded cells as quantified by the DNA assay. Error bars ± SD and ^*^p < 0.05 as per one-way ANOVA post-hoc multiple comparison analysis.

Histological analysis of the cultured scaffolds revealed cellular morphology and matrix deposition. Within the ACL fibroblast-osteoblast co-culture scaffolds, ACL fibroblasts showed higher proteoglycan deposition compared to the osteoblasts as indicated by safranin O/fast green staining (Figure 4B I, C I). Positive stains for non-crosslinked collagen type I (Figure 4B II, CII) were observed, indicating high cell-driven matrix remodelling. Tenomodulin, a mature ligament protein, was present in the ACL fibroblast phase (Figure 4B III), but not in the osteoblast phase (Figure 4C III). Scleraxis is a transcription factor that is activated for ligament and tendon maturation [31], but was detected in both fibroblast and osteoblast zones. Previously, immunohistochemical studies have indicated similar levels of scleraxis in isolated human tendon fibroblasts and osteoblasts [32]. Osteocalcin, an extracellular matrix protein produced by osteoblasts [33], was only positively stained in the osteoblast-phase. These trends within cell-driven matrix formation were also observed in single-phase ADCs that were cultured for 14 days (Figure S1). From the immunohistological stains of the co-culture, especially of collagen type I, the cell morphology could also be interpreted (Figure 4B II, C II). The ACL fibroblasts were significantly more polarized and elongated, whereas the osteoblasts displayed more randomly oriented and rounded morphology. These results suggested that both fibroblasts and osteoblasts seeded in the co-culture ADC model maintained their phenotype and morphology throughout the 14-day culture period.

**Figure 4.**
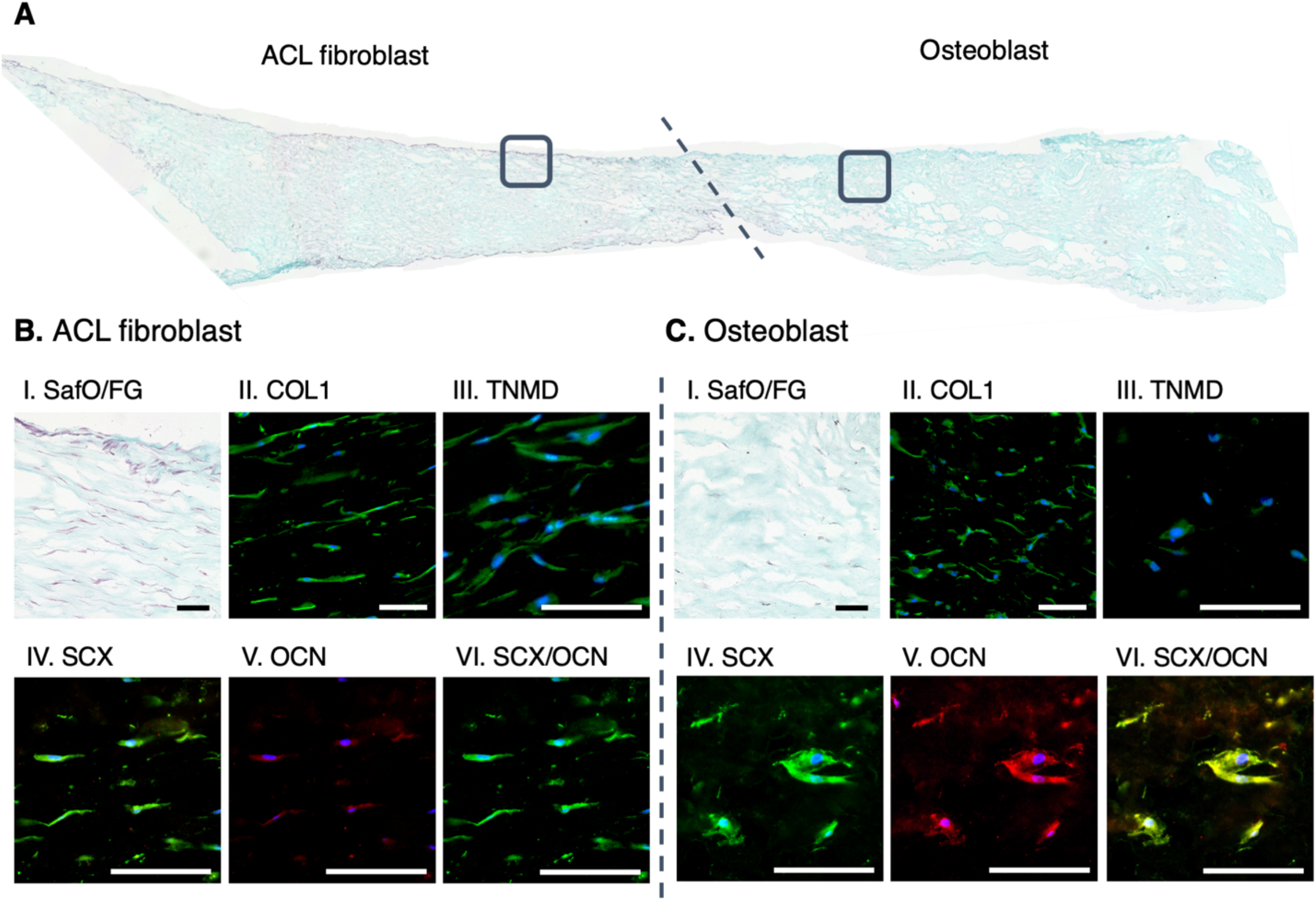
Representative images of histological and immunofluorescent analyses of two-phase ADCs after 14 days culture. (A) Safranin O/fast green (SafO/FG) of the entire construct with the ACL fibroblast and osteoblast phases separated by the dotted line. (B) ACL fibroblast phase and (C) osteoblast phase stained for (I) SafO/FG of the boxed areas of (A), (II) collagen type I (COL1), (III) tenomodulin (TNMD), (IV) scleraxis (SCX), (V) osteocalcin (OCN) and (VI) merged SCX and OCN. All scale bars = 100 µm.

## 4. Discussion

This study has demonstrated the potential application of ADC hydrogels as a robust 3D *in vitro* model of primary human ACL fibroblasts and osteoblasts within a continuous two-phase system. ADC hydrogels can be rapidly fabricated by the GAE method. Here a simple modification to the method, with two-layer casting of precursor highly hydrated collagen gels, can enable the fabrication of two-phase ADCs. Attributable to the relative viscous nature of the neutralized collagen solutions, the cells maintain their location during the 30-minute gelation period. Two neutralized collagen solutions were prepared and seeded with either fibroblasts or osteoblasts, which were layered within the gelation mould to undergo fibrillogenesis. The precursor gels were subjected to the GAE method to simultaneously compact and align the collagen fibrils [23,24], while maintaining the two-phases that were then cultured for 14 days. The anisotropic nature of ADCs resembled the structure of native ligament tissues and showed comparable mechanical properties to immature ligament, whose tensile modulus ranges between 1-3 MPa [34] and seeded mesenchymal stem cells was observed to undergo tenogenesis [26]. These ADCs have also been identified as osteoid-mimicking scaffolds [21], and have been shown to form ectopic bone *in vivo* [35]. Although the ADCs are limited in their mechanical properties when compared to mature musculoskeletal tissues, the structure, composition, and modulus mimic those of immature tissues, rendering it an appropriate scaffold material for this enthesis model.

Although ACL ligament and enthesis injury and replacement surgeries are commonplace, there is not yet a robust *in vitro* model for studying these injuries. Currently, *in vivo* models remain the gold standard [9], but these are either small and limited in their resemblance to human healing or are large and are accompanied by high costs [36]. Although *in vitro* models show less complexity, single-culture 3D models remain of interest to improve their mimicry of native musculoskeletal tissues in both structure and composition [37]. With co-culture of differentiated primary cells, a simplified representation of native tissue physiology may be achieved. In 2D fibroblast-osteoblast co-culture systems, crosstalk between the two cell types and increased fibrocartilage phenotypes have been observed [14]. Although 2D models can give insight into the cellular interactions, there is a need for 3D models that more accurately recreate the native extracellular matrix and physiological microenvironment. Collagen models that are commonly used are highly hydrated and are mechanically weak [12]. Previous investigations on collagen hydrogels seeded with high-density bovine tenocytes and ligament fibroblasts induced cell-driven matrix remodelling that reached comparable mechanical properties of immature native tendon and ligament after 4-6 weeks of culture [37]. Primary bovine cells have previously been shown to proliferate more rapidly and deposit more matrix than primary human cells, adding a layer of complexity to human translation [38,39]. As GAE fabricates the ADCs with comparable mechanical properties within 2 hours, this method significantly minimizes the need for prolonged cell culture.

Previously, one study utilised the GAE method to fabricate and investigate a 3D co-culturing system designed to model the interaction between implanted Schwann cell-seeded ADCs and native neuronal cells, *in vivo* [25]. Specifically, Schwann cells were initially seeded within the ADCs and neuronal cells were subsequently seeded on the hydrogel surface, thus highlighting the increased applications of the ADC model. Dense collagen systems not only benefit from ease of manipulation and versatility, but also from the ability to withstand mechanical stimulation without damaging the material while having a positive influence on cellular activity. Since musculoskeletal tissues, including enthesis, ligament and bone, are load-bearing and force-transmitting tissues, *in vitro* models should also enable the study of cellular mechanotransduction. It is widely accepted that mechanical loading impacts the development and healing of the enthesis [11,40–42], which has generated increasing interest in the mechanobiological effects on the enthesis and its connecting tissues [42]. Current models have yet to investigate the effects of mechanical loading to a co-culture system. The proposed 3D co-culture model is compliant up to approximately 30% strain and may be able to withstand external mechanical loading. Therefore, this *in vitro* model may be used to complement currently applied *in vivo* models.

The recent automation of the GAE method may also increase the utility of this model as a high throughput fabrication of predetermined ADC characteristics have been achieved by manipulating the casting cross-sectional area and the needle cross-sectional area [43]. Additionally, this *in vitro* model could provide a means of investigating patient-derived cells for insight into healing and personalized medicine. In this approach, cells from damaged or diseased tissues can be seeded into ADCs and stimulated either chemically or mechanically to observe potential healing mechanisms that could be harnessed for a novel therapeutic. As ligament and tendon replacement surgeries are commonplace and patient-specific models using these tissues could be fabricated to investigate variations in healing linked to variables including severity of injury, gender, age, and exercise levels. The model can also be adapted to investigate other intercellular interactions. As demonstrated in this study, a two-phase model of lung fibroblast and cancerous cell lines was fabricated as a proof-of-concept to observe cellular distributions, but could be used further to investigate cell migration or drug sensitivity. These models may be useful tools in investigating cellular crosstalk and their responses to external stimuli.

## 5. Conclusion

This work demonstrated the rapid fabrication of two-phase 3D co-culture systems. The limitations of either 2D or traditional highly hydrated collagen gels can be overcome by the densification and fibrillar realignment during the GAE fabrication process to generate ADC gels. Both acellular and cellular two-phase systems were fabricated, and ACL fibroblast-osteoblasts in ADCs maintained their phenotypes throughout 14 days. These results indicate the potential use of this two-phase scaffold as an *in vitro* model of interfacial tissues including the enthesis that can be stimulated both chemically and mechanically to further understand the mechanisms of these cells and their intercellular crosstalk.

## Supporting information

SI_control

## 7. Acknowledgements

The authors gratefully acknowledge funding from Canada NSERC, FRQNT, CFI, Research Institute McGill University Health Center start-up funds and Réseau de recherche en santé buccodentaire et osseuse (RSBO). DHR is a FRQS Junior 2 Research Scholar and HP’s funding was partially supported by McGill Engineering Doctoral Award and the Lorne Trottier Engineering Graduate Fellowship. MEC acknowledges support from the Canadian Institutes for Health Research (CIHR Funding Reference No. 171258). The authors would also like to thank McGill Scoliosis and Spine Group, McGill orthopaedic residents and Transplant Quebec for their assistance in recovery of spine samples.

