## Supplementary material for "Continuous two-phase *in vitro* co-culture model of the enthesis": SI_control

**Supplementary information**

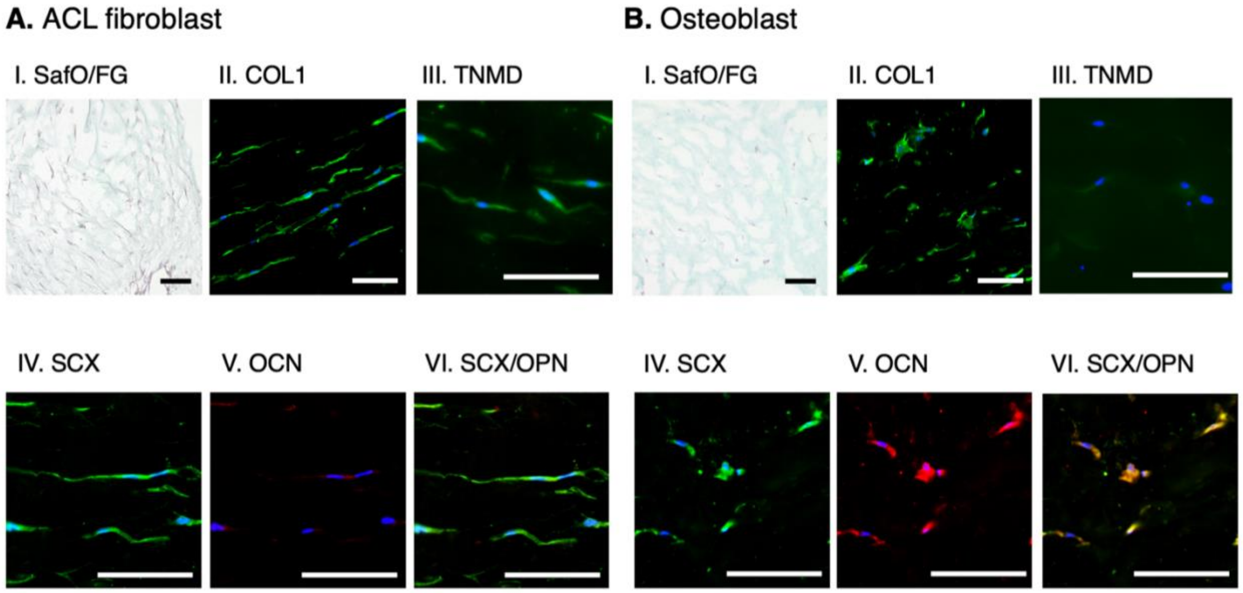


Figure S1 Histological and immunofluorescent analysis of single-phase ADCs after 14 days culture. (A) ACL fibroblast seeded ADCs and (B) osteoblast seeded ADCs stained for (I) Safranin O/fast green (SafO/FG), (II) collagen type I (COL1), (III) tenomodulin (TNMD), (IV) scleraxis (SCX), (V) osteocalcin (OCN) and (VI) merged SCX and OCN. All scale bars = 100 μm.
